# Could instructor talk drive CURE effectiveness? A comparative study of instructor talk in introductory lab courses

**DOI:** 10.1101/2025.05.09.653137

**Authors:** Christopher James Zajic, Kelly Subramanian, Arnav Adulla, Zarae Allen, Meghan B. Blitchington, Avery Brotzman, Edward Carrillo, Jheel Dhruv, Taliyah Evans, Sabrina Haider, Sehee Ashley Han, Leighton Hilton, Katie Holliday, Ethan Keairnes, Joon Kum, William Nathan Lantz, Joel Martin, Matthew Pierce-Tomlin, Mallory J. Plunkett, Cheryl S. Sam, Fama Sarr, Amanda Shroyer, Caitlin Tucker, Kenneth Walton, Madison A. West, Elizabeth Wolfson, Brandon Yoon, Koleas Zumbro, Erin L. Dolan

**Affiliations:** Department of Biochemistry & Molecular Biology, University of Georgia, Athens GA 30602; University of California Davis, Davis, CA 95616; University of Texas El Paso, El Paso, TX; Haverford College, Haverford, PA, 19041; University of North Carolina at Chapel Hill, Chapel Hill, NC 28223

**Keywords:** CURE, Lab instruction Instructor talk Immediacy, Expectancy value theory Self-efficacy, Task values

## Abstract

Course-based undergraduate research experiences (CUREs) are thought to enhance students’ motivation to continue in college, in science, and in research. Yet, how CUREs enhance student motivation is largely undefined. Theories of instructor immediacy, self-efficacy, and task values suggest that CURE instructors may talk in ways that influence students’ motivational beliefs. We characterized the non-content related talk of instructors teaching 48 introductory biology lab courses, half CUREs and half non-CUREs. We identified 14 types of instructor talk that fit these theoretical perspectives: fostering students’ closeness with their instructor (i.e., immediacy talk), building students’ confidence in their scientific abilities (i.e., self-efficacy talk), and promoting students’ sense of worth in their work (i.e., task value talk). Course type had a medium effect on talk type, with CURE instructors utilizing more immediacy, self-efficacy, and task values talk than non-CURE instructors but also showing more variation in these types of talk. Our results suggest that motivation-related instructor talk is more prevalent in CUREs than non-CUREs, but wide variation in CURE instructor talk indicates additional investigation is needed before non-content talk can be considered a mechanism for the motivational influences of CUREs.

**HIGHLIGHT:** This study compares non-content instructor talk in CURE and non-CURE lab courses using immediacy, self-efficacy, and task value theories. CURE instructors use more talk than non-CURE instructors, but variation in CURE instructor talk leaves open the question of whether talk is a causal factor in the motivational influence of CURE instruction.

## INTRODUCTION

Course-based undergraduate research experiences (CUREs) engage students in working on a research question or problem with unknown outcomes that are of interest to stakeholders outside the classroom (Auchincloss et al., 2014; Cooper et al., 2019). CUREs have grown in popularity for their potential to scale up access to undergraduate research and influence undergraduates’ retention in the sciences (Buchanan & Fisher, 2022; Gentile et al., 2017; Hanauer et al., 2017; Rodenbusch et al., 2016; Watts & Rodriguez, 2023). However, the features of CUREs that influence students’ motivation, meaning their goal-directed behavior evidenced by their persistence and success in science, remain largely unknown (Gentile et al., 2017; Schunk et al., 2014). Prior research has hypothesized features of CUREs that distinguish them from traditional lab courses, which typically engage students in well-defined lab exercises with predictable outcomes illustrating established scientific principles (Corwin, Runyon, Robinson, & Dolan, 2015; Hanauer & Dolan, 2014; Hanauer, Frederick, Fotinakes, & Strobel, 2012). Specifically, CUREs are thought to offer opportunities for students to:

- Make *discoveries*, meaning that their work has the potential to yield novel products and results that are of interest to stakeholders outside of the course;
- Engage in *iterative* work, including revising work based on feedback, collecting additional data, and repeating work to troubleshoot or problem-solve or to confirm findings; and
- Develop greater *ownership* of their work than students in traditional lab courses, which positively relates to their intentions to continue in science (Corwin et al., 2018).

Yet, these features explain only a fraction of the increase in students’ intentions to pursue a research career after participating in a CURE (Corwin et al., 2018). In addition, reviews of research on CUREs reveals that CUREs vary widely in the extent to which students have opportunities to make discoveries, engage in iterative work, and develop project ownership (Beck et al., 2023; Watts & Rodriguez, 2023). Research is needed to understand the features of CURE instruction that may shape students’ decisions to continue in science and research.

One study comparing a CURE and a non-CURE version of a lab course suggests that how the instructor frames the course – their talk – may be a critical factor (Cooper et al., 2019). In this study, students in CURE versus non-CURE versions of the course experienced the same curriculum taught by the same instructor. The only difference was that CURE students were told they were studying samples that had not previously been characterized and thus they had the opportunity to make broadly relevant discoveries.

Non-CURE students were told they were studying samples that had previously been characterized to confirm what was already known. Students in the CURE version reported significantly higher levels of discovery and ownership, but not iteration, compared with students in the non-CURE version. This study did not link discovery or ownership to student outcomes such as learning or motivational beliefs. Yet, it suggests that what instructors say during CURE instruction may influence whether and to what extent students experience these courses as motivating environments.

Instructor talk, or things instructors say aloud to students during a course, has been studied in K-12 settings for decades because of its potential to influence what students do in class, what students learn, and whether students are motivated to continue in a field of study or opt out of it (Boden et al., 2019; Nystrand, 2006; Nystrand & Gamoran, 1991; Scott, 1998; Wells, 1999; Wells & Arauz, 2006; Zepeda et al., 2019). Research at the undergraduate level has focused on “non-content instructor talk,” or what instructors say *beyond* the content of the course (Harrison et al., 2019; Seidel et al., 2015). The first study of instructor talk in undergraduate biology education defined “instructor talk” as any language that was spoken by the instructor to the whole class and was not specific to course content, content analogies, or course logistics (Seidel et al., 2015). This analysis took a “grounded approach,” ascribing qualitative meanings to instructor quotes and identifying emergent themes without a priori theoretical framing. This work identified 17 types of talk that fit five themes: building the instructor/student relationship, establishing classroom culture, explaining pedagogical choices, sharing personal experiences, and unmasking science. Subsequent research has elaborated on this initial characterization by testing its applicability to a larger and more diverse group of biology lecture instructors and identifying negative forms of instructor talk (Harrison et al., 2019).

Although this foundational work on non-content instructor talk, which we will refer to as “instructor talk” from here forward, has been important for describing talk in biology lecture courses, adaptations are needed to study lab instruction. Lab instructors regularly circle the room and speak to small groups of students, potentially influencing students’ experiences and outcomes without speaking to the whole class. Gelinas and colleagues (2022) address this difference in lab course format in their study of instructor talk observed among graduate teaching assistants (GTAs) teaching biology lab courses. They characterized the non-content language of GTAs without reference to whether GTAs were speaking to the entire class or to individual or small groups of students (Gelinas et al., 2022). Their results showed that GTAs differed in their instructor talk based on their teaching and professional development experience, demonstrating the utility of the framework for distinguishing among lab course instructors in meaningful ways. However, this research did not offer or test theory that could explain how instructor talk influences students.

Ovid and colleagues (2021) took a step toward connecting the instructor talk framework (Harrison et al., 2019; Seidel et al., 2015) with student experiences by asking students about what instructor talk they remember and whether they perceive provided examples of talk positively or negatively. This study framed positive and negative learning environments broadly. For example, students might perceive a learning environment as positive if it promotes their engagement, learning, performance, sense of belonging, persistence, and/or success. Students might perceive a learning environment as negative if they perceive limitations or undermining of any of these facets of their learning experience. Clear and coherent theory is needed to bridge the gap between systematic descriptions of instructor talk and how talk influences the experiences and outcomes of students in biology courses.

Research on non-content instructor talk outside of undergraduate biology courses has been grounded primarily in theory related to verbal instructor immediacy, or what instructors say to foster and support students’ development of a sense of closeness with their instructors (Mehrabian, 1981). Notably, Ovid and colleagues (2021) found that the majority of students in their study remembered talk that fits this theory. Other research has made theoretical connections between verbal instructor immediacy and student motivation. For example, instructor verbal immediacy, such as using humor or sharing personal information (Gorham, 1988), increases students’ learning self-efficacy and other predictors of motivated behavior (Barahona Guerrero, 2017; Velez & Cano, 2008, 2012; Witt et al., 2004). This connection raises a question of whether non-content instructor talk might explain the motivational influence that CUREs have on students. To test this idea, we sought to characterize and compare the talk used by instructors teaching introductory biology lab courses with a research component (CURE) or with no research component (non-CUREs).

We were interested in exploring instructor talk from a motivational perspective, given prior research showing CURE instruction influences students’ persistence in science (Hanauer et al., 2017; Rodenbusch et al., 2016). Thus, we focused on identifying and characterizing forms of non-content talk that had the potential to be perceived by students as motivating. To accomplish this, we used theoretical perspectives of instructor immediacy, self-efficacy, and task values to analyze the talk we observed and compare occurrences of each type of talk between CURE and non-CURE courses. This approach necessarily differs from the grounded approach of prior research on instructor talk (Gelinas et al., 2022; Harrison et al., 2019; Ovid et al., 2021; Seidel et al., 2015) because it aims to identify forms of talk related to specific constructs rather than letting the data alone drive characterization of instructor talk (Cho & Lee, 2014; Pfeifer & Dolan, 2023; Starks & Brown Trinidad, 2007; Suddaby, 2006). Using instructor immediacy as a frame allowed us to identify forms of talk that could foster a feeling of closeness between instructors and students that is thought to promote student learning and success. Using self-efficacy and task values as frames allowed us to elucidate the forms of instructor talk that could promote students’ confidence in their scientific abilities (i.e., self-efficacy) and their perceptions that doing science is worthwhile (i.e., task values). Next, we describe these perspectives and elaborate on their relevance to the study of instructor talk in CUREs.

### THEORETICAL FRAMING

Here we describe prior research related to instructor immediacy and its effects on student motivation, theorizing about how CURE and non-CURE instructors might talk in common and distinct ways that promote a sense of closeness with students. We also describe self-efficacy and task values, which are key constructs in the expectancy value theory of motivation (Wigfield & Eccles, 2000), and we theorize about how CURE versus non-CURE instructor talk might relate to these constructs.

### Immediacy

Scholars of instructor immediacy have argued that student learning can be improved when students have closer and more positive relationships with their instructors (Richmond et al., 2017). Such relationships can be fostered through verbal and non-verbal instructor behaviors to create a sense of closeness between students and their instructor. Two theoretical mechanisms have been proposed to explain the influence of instructor immediacy on students. The first is arousal attention, which posits that instructor immediacy increases students’ alertness, heightening their focus and germane effort and subsequently their learning and memory (Allen et al., 2006; Christophel & Gorham, 1995; Kelley & Gorham, 1988). The second presupposes that students learn when they want to (i.e., trait motivation) and instructor immediacy can create circumstances that increase students’ desire to learn (i.e., state motivation) (Gorham & Christophel, 1990; Richmond, 1990). For instance, students’ liking of their instructor fosters their liking of the course and the subject matter, ultimately increasing their desire to learn (Richmond et al., 2017).

Primary and meta-analytic research has shown that instructor immediacy behaviors are positively associated with motivation to learn and with learning (Comadena et al., 2007; Pogue & Ahyun, 2006; Richmond, 1990; Witt et al., 2004). However, most of this research relies on student reports of nonverbal behaviors such as instructors gesturing when talking to the class, facing the class, or moving from behind a podium and around the classroom. Verbal immediacy behaviors, such as calling students by name, interacting with students before and after class, praising students’ work, and discussing outside-of-class experiences are also thought to promote a sense of closeness. Meta-analytic research indicates that instructors’ verbal immediacy behaviors are associated with student learning and motivation, although the effects are heterogeneous (Liu, 2021; Witt et al., 2004). Investigations of instructor talk in undergraduate courses have included verbal immediacy behaviors as well as forms of instructor talk hypothesized to decrease student resistance to teaching strategies and students’ sense of stereotype threat (Harrison et al., 2019; Seidel et al., 2015).

Because CUREs are designed to engage students in research, which is substantively different from traditional lab exercises designed solely to build students’ disciplinary content knowledge and skills (Hanauer et al., 2012), CUREs may afford unique opportunities for instructors to display verbal immediacy behaviors. For example, CURE instructors may describe their own experiences with research outside of the class, while non-CURE instructors may not, given the irrelevance of traditional lab course material to their research experience. In addition, biological research often requires waiting for samples to incubate, equipment to run, or organisms to grow. Thus, CUREs may afford more opportunities for informal conversation that builds closeness when compared to traditional lab exercises, which have often been fine-tuned to be completed efficiently in a designated lab period. When instructors foster a sense of closeness with these forms of talk, students may pay more attention in class or see their instructor as more likeable. As a result, they may choose to put forth more effort that results in more success such that they ultimately choose to continue in a science or research-related path. In addition, instructors who foster closeness with verbal immediacy may build a sense of trust among students such that they are more likely to believe their instructor’s other motivation-related messages. Some of these examples overlap with the types of instructor talk described by Seidel and colleagues (2015) and in follow-on work (Harrison et al., 2019; Ovid et al., 2021), such as building the instructor/student relationship and sharing personal experiences.

### Self-Efficacy

Students’ development of scientific self-efficacy, or confidence in their ability to carry out science practices and succeed in science, is one of the most consistent and robust outcomes of CURE instruction (e.g., Dunbar-Wallis et al., 2024; Newell & Ulrich, 2022; Wilczek et al., 2022). Thus, self-efficacy theory and research on sources of self-efficacy have the potential to be useful for characterizing instructor talk in CUREs versus non-CURE lab courses. Bandura (1997) described self-efficacy as the belief that one can plan and execute actions to achieve a particular outcome. Bandura (1997) theorized and others have described four main sources of self-efficacy (Chen & Usher, 2013; Usher & Pajares, 2008). First, mastery experiences – or the successful completion of a task or accomplishment of a goal – is considered the main driver of self-efficacy. Second, vicarious experiences – or observing similar others successfully completing a task or accomplishing a goal – can drive self-efficacy growth by instilling the belief that people “like me” can be successful. Third, social persuasion – or receiving positive appraisals from respected others – can foster self-efficacy growth by reinforcing the belief that a person whose opinion is credible is gauging one’s ability favorably. Finally, positive emotional arousal like excitement or interest can promote self-efficacy and negative emotional arousal like fear or anxiety can undermine self-efficacy.

As mentioned previously, CUREs are designed to engage students in research, which may afford unique opportunities for instructors to use self-efficacy talk. CURE instructors may be better positioned to promote students’ sense of scientific mastery because they can speak to how students are making progress toward achieving a larger scientific goal (Auchincloss et al., 2014; Corwin, Graham, et al., 2015; Dolan & Weaver, 2021). When research does not proceed as anticipated, CURE instructors may encourage students to repeat techniques or experiments such that students gain more experience that builds their scientific self-efficacy (Corwin, Graham, et al., 2015; Dolan & Weaver, 2021). CURE instructors may also prompt students to trouble-shoot or problem-solve, setting a verbal expectation that students are working on a challenging task in which they will ultimately be successful given time and effort. In contrast, when students experience challenges, setbacks, or failures with traditional lab exercises, instructors may encourage them to use classmates’ data or move on to the next lab exercise. This type of talk may miss the opportunity to frame scientific challenges as a mastery experience.

CURE instructors may function as a source of social persuasion by attributing students’ scientific struggles and failures to the circumstances of research (Gin et al., 2018; Goodwin, Cary, et al., 2021). When students fail to generate expected results from traditional lab exercises, non-CURE instructors may attribute these results to “human error,” potentially persuading students that they are not scientifically capable (Corwin & Charkoudian, 2022). Because students typically work together in lab courses, both CUREs and non-CUREs may foster vicarious experiences by affording opportunities to observe peers’ successes. Yet, students may perceive peers’ successes in the context of a CURE as an example of how to be successful in research, not just in lab class. Instructors may foster these vicarious experiences by talking to students about working together or learning from one another (Auchincloss et al., 2014; Beck et al., 2023; Brownell & Kloser, 2015). Finally, CURE instructors may reduce students’ negative emotions by normalizing the challenges and failures associated with doing research (Gin et al., 2018; Goodwin, Anokhin, et al., 2021; Henry et al., 2019). Some of these examples overlap with the types of talk described in the general instructor talk framework (Harrison et al., 2019; Ovid et al., 2021; Seidel et al., 2015), such as boosting self-efficacy, building a community among students, and indicating it is ok to be wrong or to disagree.

### Task Values

The expectancy value theory of motivation couples competency-related beliefs – meaning beliefs in one’s abilities related to a task or goal (i.e., expectancies) – with value beliefs, or whether a task or goal is worthwhile to pursue, to explain motivated behavior (Eccles & Wigfield, 2002, 2020; Wigfield & Eccles, 2000). Researchers have theorized and empirically demonstrated four main components of task value (Gaspard, Dicke, Flunger, Brisson, et al., 2015; Gaspard, Dicke, Flunger, Schreier, et al., 2015; Harackiewicz & Hulleman, 2010; Wigfield & Eccles, 1992). First is intrinsic value, or the enjoyment or interest associated with doing a task. Second is attainment value, or the importance associated with doing well on the task or achieving the goal. Third is utility value, or how useful the task or goal is to accomplish. Utility value includes various forms of usefulness, including utility for daily life, academic success, career prospects, or community impacts. The final component is cost, or the negative consequences associated with the task or goal. Costs include actual financial costs, emotional costs, and opportunity costs, meaning what opportunities may be sacrificed to complete the task or accomplish the goal (Barron & Hulleman, 2015; Flake et al., 2015).

CUREs may function as a distinctively favorable environment for instructors to emphasize the value of students’ lab coursework. Research on why faculty choose to teach CUREs reveals that they find CURE instruction more intellectually stimulating and professionally rewarding (Shortlidge et al., 2017). Thus, CURE instructors may foster students’ perceptions of the intrinsic value of their lab coursework by talking about their own interest in and enjoyment of the work in a way that non-CURE instructors do not. CURE instructors may promote students’ beliefs about the attainment value of their coursework by talking about its relevance beyond the classroom and the potential to contribute to a larger body of knowledge (Auchincloss et al., 2014; Shortlidge et al., 2017). While both CURE and non-CURE instructors may influence students’ beliefs about the utility of their coursework by talking about how the knowledge and skills they develop may be useful in other contexts or in the future, only CURE instructors can reference how students’ scientific results could make a difference in the world (Boucher et al., 2017; Dunbar-Wallis et al., 2024; Fuhrmeister et al., 2021; Hanauer et al., 2017). Finally, CURE instructors may mitigate student perceptions of the costs associated with coursework by noting that scientific failures will not result in failing grades or that the effort invested in the course will “pay off” in the end (Dolan & Weaver, 2021). Again, some of these examples map onto types of talk described in the general instructor talk framework (Harrison et al., 2019a; Ovid et al., 2021; Seidel et al., 2015), such as connecting biology to the real world and career, fostering wonder in science, and being explicit about the nature of science.

### The Current Study

We used theoretical frames of immediacy, self-efficacy, and task values to characterize and compare the instructor talk we observed in CURE vs. non-CURE lab courses. Specifically, we collected multiple audio recordings from a cross-institutional sample of instructors teaching 48 lab courses. We transcribed the recordings and conducted a combination of deductive and inductive qualitative content analysis using these three theoretical perspectives, drawing from prior categorizations of non-content instructor talk when relevant to each perspective. We then tested whether immediacy, self-efficacy, and task values talk differed between CURE and non-CURE courses.

## METHODS

The current study involved collecting and analyzing two types of data from two groups of participants: (1) audio recordings of instructors as they taught lab courses, which we used to characterize their non-content talk; and (2) student survey responses on their perceptions that their courses had features associated with CURE instruction, which we used to independently confirm instructor descriptions of their courses as including a research component. The results reported here are part of a larger study that was reviewed and determined to be exempt by the University of Georgia Institutional Review Board (PROJECT00003103).

### Non-Content Instructor Talk

To characterize types of non-content instructor talk in CUREs versus non-CUREs, we collected class recordings from a North American sample of 48 instructors teaching introductory life science lab courses (Table 1). An additional 12 instructors who started the study were not included in the final analytic sample due to a lack of submitted recordings or insufficient student participation. To determine whether instructors were teaching a CURE or a non-CURE, we asked instructors how many weeks there were in their course and, of this total, how many weeks they spent on each of the following components:

- **Research:** Students worked on a research question or problem with unknown outcomes that are of interest to stakeholders outside the classroom;
- **Inquiry:** Students developed and carried out their own experiments or projects to address a question or problem that is interesting to them but not relevant to stakeholders outside the classroom;
- **Concepts:** Students completed an established set of activities that demonstrate well-understood scientific principles; and/or
- **Skills:** Students completed an established set of techniques in order to develop technical skills.

**Table 1.** Instructor demographics. Percentages reflect proportion within the total sample or each sub-sample (i.e., within CUREs or non-CUREs).

|  | **Total Sample** | **CURE Sample** | **Non-CURE Sample** |
| --- | --- | --- | --- |
|  | 48 | 24 | 24 |
| **Sex** |  |  |  |
| Female | 35 (73%) | 16 (67%) | 20 (83%) |
| Male | 10 (21%) | 7 (30%) | 3 (13%) |
| Other | 3 (6%) | 1 (4%) | 1 (4%) |
| **Race/Ethnicity** |  |  |  |
| Asian | 6 (13%) | 3 (13%) | 2 (8%) |
| Black or African American | 4 (8%) | 3 (13%) | 1 (4%) |
| Hispanic or Latinx | 3 (6%) | 1 (4%) | 2 (8%) |
| White | 38 (79%) | 19 (79%) | 19 (79%) |
| Other | 2 (4%) | 0 | 2 (8%) |
| **Position** |  |  |  |
| Graduate Student | 8 (17%) | 4 (17%) | 4 (17%) |
| Postdoctoral Position | 1 (2%) | 1 (4%) | 0 |
| Lecturer | 6 (13%) | 2 (8%) | 4 (17%) |
| Assistant Professor | 10(21%) | 6 (25%) | 4 (17%) |
| Tenured Professor | 12 (25%) | 6 (25%) | 6 (25%) |
| Other | 11 (23%) | 5 (21%) | 6 (25%) |
| **Teaching Experience (Years)** |  |  |  |
| 1-4 | 7 (15%) | 5 (10%) | 2 (4%) |
| 5-9 | 17 (35%) | 8 (17%) | 9 (19%) |
| 10+ | 24 (50%) | 11 (46%) | 13 (54%) |
| **Institutional Setting** |  |  |  |
| Community colleges | 6 (13%) | 0 | 6 (25%) |
| Liberal arts colleges | 3 (6%) | 1 (4%) | 2 (8%) |
| Comprehensive universities | 12 (25%) | 6 (25%) | 6 (25%) |
| Doctoral universities | 27 (56%) | 17 (71%) | 10 (42%) |

We considered a course a CURE if the instructor indicated their students conducted at least one week of research. Courses with no research weeks were considered non-CUREs. Within our courses containing a research component, there was a wide variation in the total number of weeks dedicated to research (Range = 1-13 weeks, μ = 6.09, SD = 3.22) and the percentage of the course spent doing research (Range = 7% - 100%, μ = 42%, SD = 26%).

### Recruitment

We recruited instructors by email announcements to CURE- and biology lab course-related listservs, social media posts, and direct emails to lab instructors. We also asked participants and other colleagues to share the study invitation, including consent information, with instructors who met the eligibility criteria. Instructors were eligible to participate in the study if they were teaching an introductory life science lab course that they had taught before (i.e., it wasn’t their first time teaching the course). Our intention with this selection criterion was to capture talk that had the potential to be more stable and informed by prior experience rather than talk that reflected the instructor figuring out how to teach the course. Instructors who participated in the study received a $50 gift card and an individualized report of their talk results as compensation.

### Data Collection

Instructors were asked to record four separate class sessions in their entirety during a single term. Instructors were asked to record their first day of class based on prior research indicating the importance of this class for students’ motivation and outcomes (Wilson & Wilson, 2007). Instructors were asked to select three other sessions to record, spaced across the semester and reflecting the diversity of class activities in the course. Audio recordings were collected via devices worn by the instructors, unless another means of recording was preferable (i.e., via Zoom). A total of 43 instructors submitted four recordings, four submitted three recordings, and one submitted two. All submitted their first-day-of-class recording. The total recording duration for each instructor varied widely, with a minimum of 135 minutes to a maximum of 739 minutes (M = 424 minutes, SD = 161 minutes).

### Data Analysis

We used the service Rev.com to transcribe all recordings. We then checked all transcripts for accuracy and renamed the transcript blind to the type of course being taught. Although we blinded the transcripts to reduce potential for bias in coding, it was often evident from what instructors were saying that the course was a CURE or a non-CURE. We performed qualitative content analysis to identify and define distinct forms of non-content instructor talk. We utilized a combination of inductive and deductive coding during the iterative codebook development process. Our initial codebook was informed by prior research on non-content talk and mentoring functions (Eby et al., 2013; Harrison et al., 2019; Seidel et al., 2015) given our interest in identifying forms of talk that were distinctive to research instruction (CUREs) versus other forms of lab instruction (non-CUREs). We defined “non-content talk” as anything instructors said that was unrelated to specific biological concepts or techniques and that included enough substantive words to ascribe meaning. Thus, our coding captured talk related to the nature and practices of science (e.g., “this is how science works”), but not to specific biological information, concepts, techniques, or procedures (e.g., “there is a table with the reagents for calculating what the amounts of each reagent you should be adding to your PCR mix”). We coded a quote when we interpreted it as having coherent meaning. We did not code quotes that were too brief to ascribe sufficient meaning to (e.g., “Great!”). We focused on forms of talk that we theorized would differ between CUREs and non-CUREs as described in our Theoretical Framework.

During our initial coding process, two members of the research team independently coded each transcript in its entirety (i.e., the complete recording of a class session) and discussed their coding to consensus. In some instances, a third research team member was included in the discussion to bring an additional perspective and help reach consensus. As we coded, we made notes about emergent forms of talk not being captured with existing codes or ambiguity in the definitions of existing codes. We discussed these issues with the entire research team and added to or revised the codebook as needed. The iterative process of coding, discussing, and revising continued until we reached saturation, meaning that the codebook included all forms of talk we were observing, and no new forms of talk were emerging.

Once the codebook stabilized, we coded the entire corpus using the final codebook. For this phase of coding, we coded each transcript in pairs (i.e., partner-coding) and then representatives of each pair discussed to consensus. This revised coding process allowed us to ensure multiple voices and perspectives were represented during both the coding and discussion phases. This process also ensured that each transcript was reviewed by at least four team members, helping to maintain consistency in application of the codes among coders over time and to minimize the chance that talk instances were missed in the very large corpus. Most of the researchers involved in coding were undergraduate students, all of whom had completed introductory biology lab coursework and thus provided additional perspective on how undergraduate students might perceive and understand what instructors were saying. Finally, we reviewed all codes with respect to our theoretical perspectives, using quotes to clarify which perspective best reflected the data. For example, instructors in our study used talk that emphasized their availability outside of class, which reflected instructor immediacy but not self-efficacy or task values. Thus, we grouped this code into the frame of Immediacy Talk.

To compare talk types between CUREs and non-CUREs, we summed all instances of immediacy talk, self-efficacy talk, and task values talk for each instructor. We divided this value by the total minutes of recording for that instructor and multiplied by 60 to generate three values for each instructor: immediacy talk per hour, self-efficacy talk per hour, and task values talk per hour. We then performed Mann Whitney U tests (Mann & Whitney, 1947) using the “stats” package in the R statistical software (R Core Team, 2022) to test for differences in talk between CURE and non-CURE instructors. For each of the three major forms of talk, we also conducted Kolmogorov-Smirnov (KS) tests, which is a non-parametric test that compares the distributions of each talk type in CUREs and non-CUREs.

### Course Features

We surveyed students in the courses being studied to determine the extent to which they experienced course features associated with CURE instruction.

### Recruitment

We recruited student participants by asking instructors to announce the study and distribute study invitations with consent information to students in their courses. Our final analytic sample consisted of 462 students, with at least three students from each course (Table 2). Students received a $25 gift card for their participation.

**Table 2.** Student demographics. Percentages reflect proportion within the total sample or each sub-sample (i.e., within CUREs or non-CUREs). Students had the option to select multiple races/ethnicities. Thus, percentages within these demographics may total to >100%.

|  | **Total Sample** | **CURE Sample** | **Non-CURE Sample** |
| --- | --- | --- | --- |
|  | 462 | 261 | 201 |
| **Sex** |  |  |  |
| Female | 349 (76%) | 201 (77%) | 148 (74%) |
| Male | 91 (20%) | 49 (19%) | 42 (21%) |
| Other | 10 (2%) | 6 (2%) | 4 (2%) |
| Prefer not to respond | 12 (3%) | 5 (2%) | 7 (3%) |
| **Race/Ethnicity** |  |  |  |
| American Indian or Alaskan Native | 4 (<1%) | 2 (<1%) | 2 (<1%) |
| Asian | 130 (28%) | 80 (31%) | 50 (25%) |
| Black or African American | 49 (11%) | 31 (12%) | 18 (9%) |
| Hispanic or Latinx | 76 (16%) | 52 (20%) | 24 (12%) |
| Native Hawaiian or Other Pacific Islander | 2 (<1%) | 2 (<1%) | 0 |
| White | 235 (51%) | 121 (46%) | 114 (57%) |
| Other | 11 (2%) | 7 (3%) | 4 (2%) |
| Prefer not to respond | 8 (2%) | 3 (1%) | 5 (2%) |
| **Year in College** |  |  |  |
| First Year | 262 (57%) | 177 (68%) | 85 (42%) |
| Second Year | 115 (25%) | 49 (19%) | 66 (33%) |
| Third Year | 46 (10%) | 18 (7%) | 28 (14%) |
| Fourth Year | 28 (6%) | 17 (7%) | 11 (5%) |
| Other | 8 (2%) | 0 | 8 (4%) |
| Prefer not to respond | 3 (<1%) | 0 | 3 (1%) |
| **Prior Research** |  |  |  |
| None | 173 (37%) | 107 (41%) | 66 (33%) |
| 1 Term | 146 (32%) | 85 (33%) | 61 (30%) |
| 2 Terms | 76 (16%) | 44 (17%) | 32 (16%) |
| 3 Terms | 26 (6%) | 9 (3%) | 17 (8%) |
| >3 Terms | 32 (7%) | 13 (5%) | 19 (10%) |
| Prefer not to respond | 9 (2%) | 3 (1%) | 6 (3%) |

### Data Collection

We collected survey data from students at the end of their course using the secure survey service, Qualtrics. For this study, we collected data using two scales from the Laboratory Course Assessment Survey (LCAS) (Corwin, Runyon, et al., 2015) and one scale from the Project Ownership Survey (POS) (Hanauer & Dolan, 2014). We did not survey students using the collaboration scale of the LCAS because it does not reliably distinguish between CURE and non-CURE courses (Beck et al., 2023; Corwin, Runyon, et al., 2015). We briefly describe each scale here. We provide results of confirmatory factor analyses and other information about the validity and reliability of these measures in the Supplemental Materials. These data were collected as part of a longer survey that included two items as attention checks. We included students’ responses in our final analytic sample if they completed both attention checks accurately. A total of 609 students responded to our survey; 147 were removed due to failed attention checks and incomplete responses. This resulted in a final analytic sample of 462 students.

#### Discovery

We used a 5-item scale to measure students’ perceptions that their course included opportunities to make broadly relevant discoveries, meaning novel results with potential to be of interest to stakeholders outside of the course (Corwin et al., 2015). A sample item is “In this course, I was expected to develop new arguments based on data.” Response options ranged from 1 (strongly disagree) to 5 (strongly agree).

#### Iteration

We used a 6-item scale to measure students’ perceptions that their course included opportunities to iterate upon their work (Corwin, Runyon, et al., 2015). A sample item is “In this course, I had time to change the methods of the investigation if it was not unfolding as predicted.” Response options ranged from 1 (strongly disagree) to 5 (strongly agree).

#### Cognitive Ownership

We used an 8-item scale to measure students’ perceptions that they had cognitive ownership of their coursework (Hanauer & Dolan, 2014). A sample item is “I was responsible for the scientific outcomes in this course.” Response options ranged from 1 (strongly disagree) to 7 (strongly agree). We opted not to survey students about their emotional ownership given validity concerns about the measure (Corwin, Runyon, et al., 2018).

### Data Analysis

To test for relationships between potentially distinctive features of CUREs (discovery, iteration, and cognitive ownership) and course-type (CURE vs. non-CURE), we employed linear mixed effects models (LMMs). We utilized the “lmerTest” package (Kuznetsova et al., 2017) in R statistical software (R Core Team, 2022). We fit separate models for each design feature, and our models took on this form:

*student perceptions of design feature ∽ course-type (CURE vs. non-CURE) + (1|instructor)*

We normalized values for discovery, iteration, and ownership to facilitate comparison of the effects. We included course type (CURE or non-CURE) as a fixed effect in our model by dummy coding (1=CURE, 0=non-CURE) based on whether instructors described their courses as including a research component. We included instructor as a random effect because intraclass correlations showed that student responses displayed a high degree of between-group variance. See Supplemental Material for details.

## RESULTS

From our qualitative content analysis of the 182 course recordings, totaling over 328 hours, we identified 14 unique types of instructor talk aligned with our theoretical frames of immediacy, self-efficacy, and task values. We define and describe each talk type and interpret how each relates to the corresponding frame (i.e., our finalized codebook). Within each frame, we present the types of talk in order of prevalence in the data.

### Immediacy Talk

We identified four types of instructor talk that reflected verbal immediacy behaviors: *sharing personal experiences, engaging in small talk, building trust,* and *emphasizing availability* (Table 3). Immediacy talk was observed slightly more than other talk themes in our sample, comprising a total of 716 quotes or 35% of the talk corpus.

**Table 3.** Immediacy talk. We identified four types of non-content instructor talk that reflected verbal immediacy behaviors. Each type is defined and accompanied by example quotes. Collectively, immediacy talk comprised 716 quotes, or 35% of the corpus.

| Categories | Example Quotes |
| --- | --- |
| **Sharing personal experiences:** Instructor discloses their own experiences or anecdotes or relates their own experiences to the experiences of their students. (n=206; 10%) | • “This lab holds an extra special place in my heart because it was the first lab I took here when I was a student. In fact, I sat right there.”<br>• “HIV also hit a little close to home. I had an uncle who died from HIV AIDS. He had been doing intravenous drugs. He infected himself. And this is before there was much treatment or antiretroviral therapies for HIV.” |
| **Engaging in small talk:** Instructor initiates conversation that prioritizes the interaction over the content (i.e., the topic of conversation is trivial). (n=200; 10%) | • Instructor: “Where did the pie come from?” Student: “It's a pie I made for Thanksgiving.” Instructor: “That's very professional looking.”<br>• Instructor: “Do you drink a lot of tea?” Student: “Yeah, a lot.” Instructor: “Do you drink it black?” Student: “Yeah, only black.” |
| **Building trust:** Instructor emphasizes their benevolence (i.e., they care for students) and integrity (i.e., they are a reliable source of support). (n=178; 9%) | • “I really want to emphasize I'm here to support you. If you ever are in need of resources, if you're ever not sure of where to turn for your mental or physical wellbeing, please reach out, please let me know, happy to point you the right way.”<br>• “I never want to ever make anyone in here feel terrible about themselves... I work really hard to make sure that you feel like you belong in this classroom, and I respect you as an individual...” |
| **Emphasizing availability:** Instructor explicitly offers help outside of regular class time. (n=132; 7%) | • “You can come and see me whenever you want because my doors are open.”<br>• “If you are super confused about anything, shoot me an email or come see me in office hours, and I'll just make sure you're caught up and you're good to go.” |

Two forms of immediacy talk appeared to humanize instructors or make them more relatable to students, with the potential to foster closeness. First, instructors *shared personal experiences* with their students by describing events in their own lives or their personal perspectives on coursework and other topics.

Instructors shared information that gave students insight into their personal lives such that they appeared to be whole people. For instance, one instructor described how they experienced a family medical crisis, noting that “it was a very traumatic experience dealing with someone that’s [had a traumatic injury] and I’ve never experienced that before and it’s really stressful.” Instructors conveyed they understood aspects of students’ experiences and thus could relate. For example, instructors commented on how they “know what it was like to be in [their students’] shoes” or that they “know what it’s like to be in a class like this.” Instructors also humanized themselves or made themselves more relatable by *engaging in small talk*. Instructors initiated casual conversations that aimed to promote interaction, and the specific content of the conversation was unimportant. For example, instructors often asked about students’ weekend activities or what they did during school breaks.

Instructors engaged in other forms of immediacy talk that appeared more relational in nature, including talk aimed at *building trust* and *emphasizing availability*. Regarding *building trust*, instructors explained that they could be “support systems” for students and that they wanted students to “feel like [the classroom] is [their] home.” In some cases, instructors elaborated on situations students might be facing and how they could offer help, as this instructor explains:

> If you have mental health needs, relationship problems, legitimate medical problems, you’re a student with disabilities, you need help figuring out how to get your accommodations. These links will help you figure out where you need to go. And if you’re not sure after going through this, I’d be happy to put you in the right direction.

Instructors also messaged that they could support students by *emphasizing availability* outside of scheduled class time, during which they could help students with class-related tasks. Instructors would remind their students that they are “always welcome to send [the instructor] an email” or that they “can set up a time to chat online or in person.” CURE instructors in our sample used more immediacy talk per hour (Median = 2.4 occurrences per hour) than non-CURE instructors (Median = 2.0 occurrences per hour) (Figure 1A). A Mann-Whitney U test comparing CUREs (n=24) and non-CUREs (n=24) showed a medium effect (Gignac & Szodorai, 2016; Hemphill, 2003) of course type on immediacy talk (U=493, r = 0.283, CI [0.03, 0.53], *p* = 0.051). We conducted a Kolmogorov-Smirnov (KS) test to compare the distributions of immediacy talk used by CURE and non-CURE instructors. Visual comparison (Figure 1B) reveals differences in the distributions of CURE and non-CURE instructors’ immediacy talk with the largest difference (D(48)=0.29; *p*=0.26) around 2.5 occurrences per hour. This difference indicates that we observed at least 2.5 instances of immediacy talk per hour in 50% of CURE instructors, but we observed this high of a frequency in only 25% of non-CURE instructors. Although this D value is noteworthy, this difference did not meet the traditional threshold for statistical significance.

**Figure 1.**
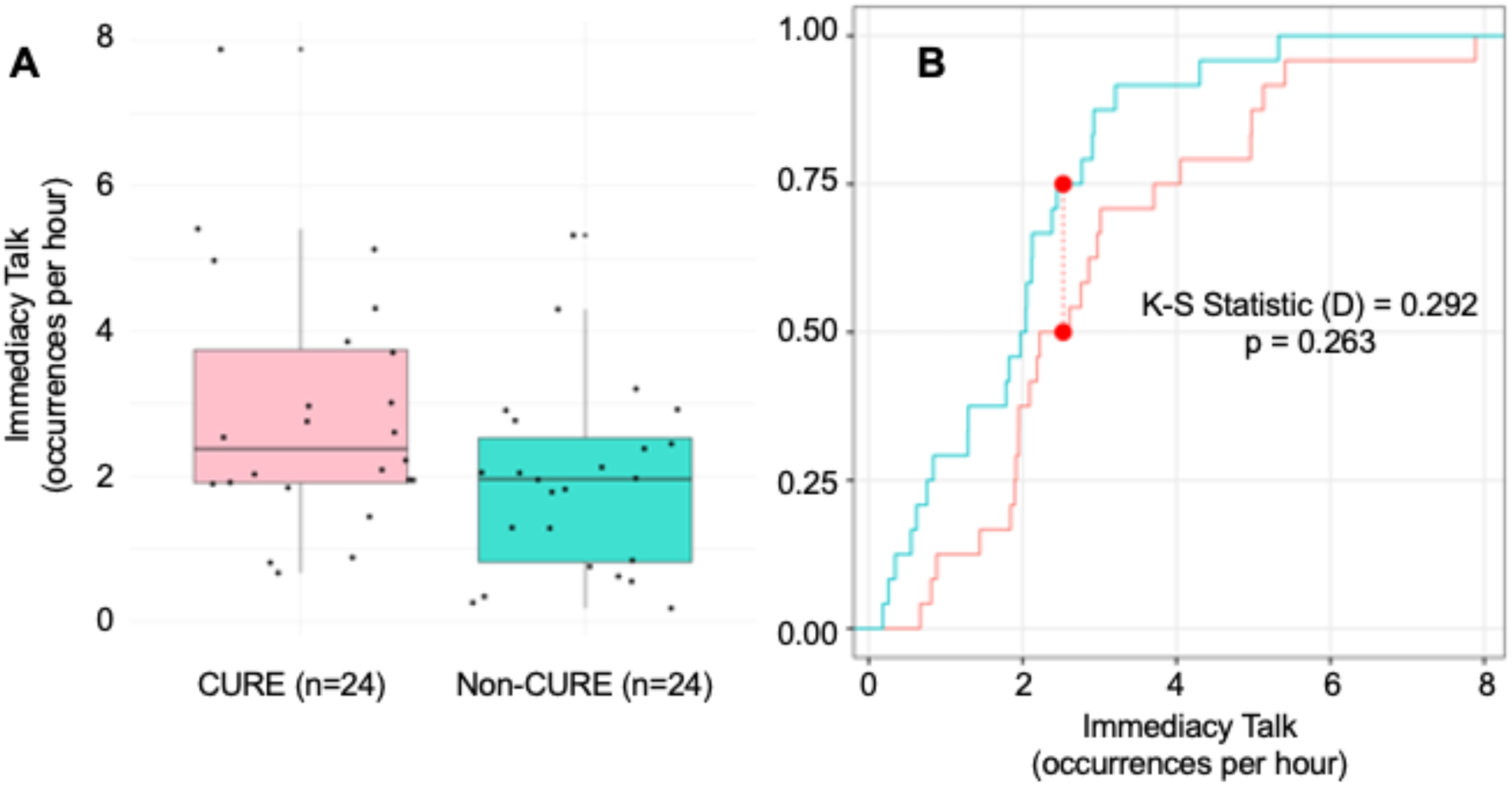
Immediacy talk in CUREs vs. non-CUREs. We compared occurrences of immediacy talk per hour for CURE instructors (coral) and non-CURE instructors (turquoise), finding that CURE instructors used more immediacy talk per hour than non-CURE instructors (U=493, r = 0.283, CI [0.03, 0.53], *p* = 0.051) (A). We also compared the distribution of immediacy talk between CURE and non-CURE instructors via a Kolmogorov-Smirnov test (B). We found that the point of greatest difference between the cumulative distribution functions (D(48)=0.29) is located between two and three occurrences per hour. ∼75% of non-CURE instructors used immediacy talk less than three times per hour, compared to ∼50% of CURE instructors. However, this difference was not significant (*p*=0.26).

### Self-Efficacy Talk

We identified six types of talk that appeared aimed at building students’ scientific self-efficacy: *normalizing struggles, providing encouragement, normalizing iteration, encouraging interdependence, encouraging independence and ownership*, and *scaffolding* (Table 4). Self-efficacy talk comprised a total of 683 quotes or 34% of the talk corpus.

**Table 4.**
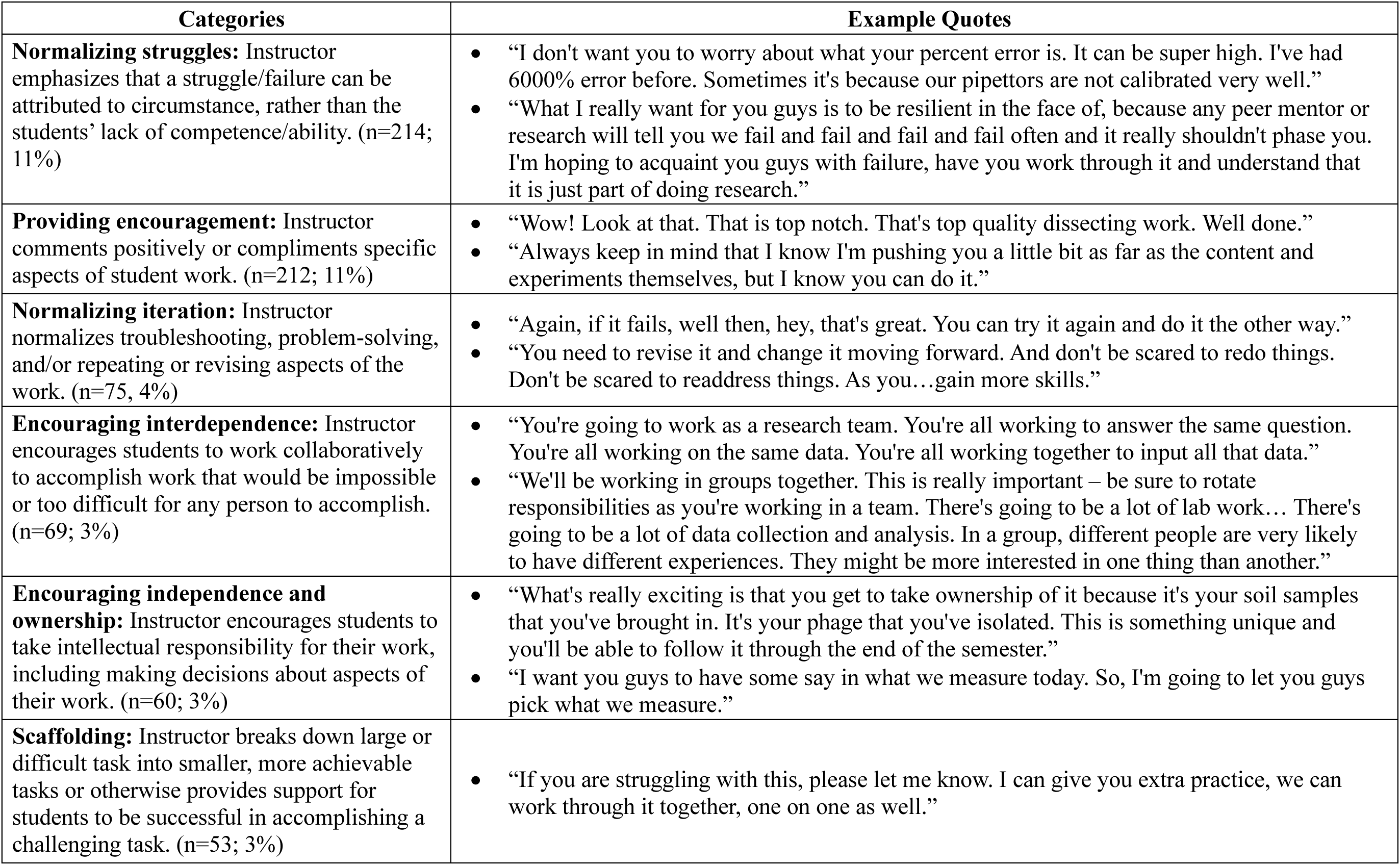
Self-efficacy talk. We identified six forms of non-content instructor talk that appeared aimed at building students’ confidence in their ability to succeed in the coursework or in science. Each form is defined and accompanied by example quotes. Collectively, self-efficacy related talk comprised 683 quotes, or 34% of the corpus.

| Categories | Example Quotes |
| --- | --- |
| **Normalizing struggles:** Instructor emphasizes that a struggle/failure can be attributed to circumstance, rather than the students' lack of competence/ability. (n=214; 11%) | • "I don't want you to worry about what your percent error is. It can be super high. I've had 6000% error before. Sometimes it's because our pipettors are not calibrated very well."<br>• "What I really want for you guys is to be resilient in the face of, because any peer mentor or research will tell you we fail and fail and fail and fail often and it really shouldn't phase you. I'm hoping to acquaint you guys with failure, have you work through it and understand that it is just part of doing research." |
| **Providing encouragement:** Instructor comments positively or compliments specific aspects of student work. (n=212; 11%) | • "Wow! Look at that. That is top notch. That's top quality dissecting work. Well done."<br>• "Always keep in mind that I know I'm pushing you a little bit as far as the content and experiments themselves, but I know you can do it." |
| **Normalizing iteration:** Instructor normalizes troubleshooting, problem-solving, and/or repeating or revising aspects of the work. (n=75, 4%) | • "Again, if it fails, well then, hey, that's great. You can try it again and do it the other way."<br>• "You need to revise it and change it moving forward. And don't be scared to redo things. Don't be scared to readdress things. As you...gain more skills." |
| **Encouraging interdependence:** Instructor encourages students to work collaboratively to accomplish work that would be impossible or too difficult for any person to accomplish. (n=69; 3%) | • "You're going to work as a research team. You're all working to answer the same question. You're all working on the same data. You're all working together to input all that data."<br>• "We'll be working in groups together. This is really important – be sure to rotate responsibilities as you're working in a team. There's going to be a lot of lab work... There's going to be a lot of data collection and analysis. In a group, different people are very likely to have different experiences. They might be more interested in one thing than another." |
| **Encouraging independence and ownership:** Instructor encourages students to take intellectual responsibility for their work, including making decisions about aspects of their work. (n=60; 3%) | • "What's really exciting is that you get to take ownership of it because it's your soil samples that you've brought in. It's your phage that you've isolated. This is something unique and you'll be able to follow it through the end of the semester."<br>• "I want you guys to have some say in what we measure today. So, I'm going to let you guys pick what we measure." |
| **Scaffolding:** Instructor breaks down large or difficult task into smaller, more achievable tasks or otherwise provides support for students to be successful in accomplishing a challenging task. (n=53; 3%) | • "If you are struggling with this, please let me know. I can give you extra practice, we can work through it together, one on one as well." |

Several types of self-efficacy talk appeared aimed at supporting students through the challenges of the scientific work they were completing in their lab courses, including both scientific practices such as collecting and analyzing data and technical procedures such as pipetting or using equipment with proficiency. Self-efficacy talk was not about providing instructions for scientific practices or guidance on how to carry out techniques or procedures, which we considered content talk. Rather, self-efficacy talk reflected what instructors said with the apparent aim of boosting students’ confidence about their scientific abilities. For example, instructors *normalized struggles* students were encountering with their scientific work, noting that issues like experimental failures and uncertain results were part of doing science rather than something students did wrong. This form of talk may protect students against blaming themselves or their own capabilities. In some cases, instructors used these occurrences as an opportunity to emphasize that students were becoming scientists. For example, one instructor elaborated that:

> It’s okay that it didn’t work. Welcome to science. I mean that in the most sincere way. Getting inconclusive or unexpected or missing results doesn’t mean that you’ve failed. It doesn’t mean you’re bad at science. It doesn’t mean you don’t belong here.

When students experienced scientific setbacks or failures, instructors also *normalized iteration*, delineating why it is scientifically valuable to repeat scientific work. For example, one instructor spoke about how iterative work was important when students generated unexpected results, explaining “This is why we have replicates. Is what happened in yours due to just mistake? Or is it actually due to the fact that they can’t survive in that high level of sugar?” By normalizing iteration, instructors may buffer against students interpreting the need to repeat work as a sign of their own shortcomings. By setting an expectation that students repeat their work, instructors may also be creating a “mastery experience” environment where students practice more and succeed at a challenging task, ultimately becoming more confident in their scientific skills.

Instructors *provided encouragement* to their students by praising their scientific progress and accomplishments. For example, one instructor complimented their students’ work, remarking, “Look at these data. They’re beautiful. I’m so thrilled. They said sometimes when you teach this lab, nobody gets the right curves, but these curves are looking great.” This kind of praise may be interpreted by students as a form of social persuasion, meaning that someone credible thinks they are capable.

Instructors *encouraged interdependence* among students, emphasizing that certain tasks would be too challenging to complete alone or that more scientific work could be completed, and the work would be more manageable if workloads were shared. For example, instructors told students to “divvy up responsibility” when they had a tedious task, like “a lot of counting.” When instructors encourage interdependence, students may view the work as more feasible than if they had to work alone or be able to make more progress as a group and feel a greater sense of accomplishment as a result. Interdependent work may also afford more opportunities to observe each other’s strategies and successes and thus feel more confident about their own capabilities.

Instructors also talked about opportunities students had for autonomy in their scientific work when they *encouraged independence and ownership*. For instance, one instructor explained to students that they were going to be “doing actual research on a question that you designed, a real experiment that you designed.” Again, students may perceive this talk as an indication they are capable of making decisions and carrying out the work rather than needing their instructor to tell them how.

When students were working on challenging tasks, instructors sometimes intervened with *scaffolding* talk, explaining how a large or challenging task would be accomplishable in smaller, more achievable chunks and how they would provide support for students to be successful. For example, one instructor acknowledged that “the bars are set high” in their class but followed with an explanation that they are “going to be supporting [students] …to help them get closer to reaching the goal.” Having indicators of progress (i.e., getting “closer”) and providing support in achieving goals may help students feel more confident in their capabilities.

CURE instructors used more self-efficacy talk (Median = 2.7 occurrences per hour) than non-CURE instructors (1.6 occurrences per hour) (Figure 2A). A Mann-Whitney U test comparing CUREs (n=24) and non-CUREs (n=24) showed a medium effect (Gignac & Szodorai, 2016; Hemphill, 2003) of course type on self-efficacy talk (U=502, r = 0.246, CI [0.02, 0.53], *p* = 0.078). Visual inspection of the results of a KS test revealed greater difference between CURE and non-CURE instructors’ use of self-efficacy talk at the high end of the distribution (Figure 2B). In other words, about half of the instructors in our study used self-efficacy talk 1-2 times per hour regardless of their course type. However, a greater proportion of CURE instructors used more self-efficacy talk per hour, with at least half the CURE instructors in our sample using >3 instances per hour and 25% of CURE instructors using 4+ instances per hour. In contrast, ∼10% of non-CURE instructors in our sample were observed using 3 instances per hour and none were observed using 4 instances per hour. The KS statistic (D(48)= 0.375) at ∼3 instances per hour supports this observation, although the difference did not meet the traditional threshold for statistical significance (*p*=0.068).

**Figure 2.**
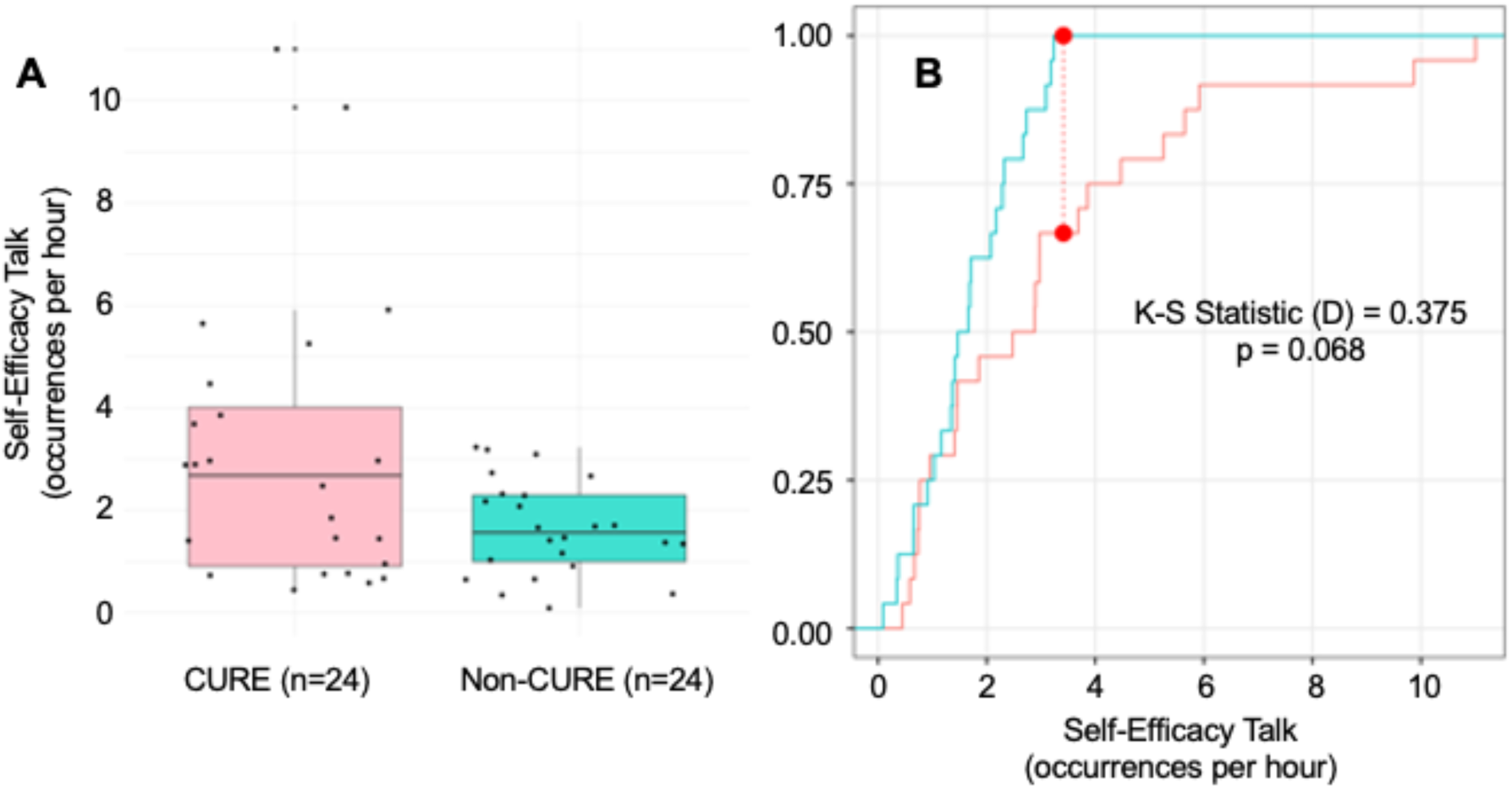
Self-efficacy talk in CUREs vs. non-CUREs. We compared occurrences of self-efficacy talk per hour for CURE instructors (coral) and non-CURE instructors (turquoise), finding that CURE instructors used more self-efficacy talk per hour than non-CURE instructors (U=502, r = 0.246, CI [0.02, 0.53], *p* = 0.078) (A). We also compared the distribution of self-efficacy talk between CURE and non-CURE instructors via a Kolmogorov-Smirnov test (B). We found that the point of greatest difference between the cumulative distribution functions (D(48)= 0.375) is located between three and four occurrences per hour. 100% of non-CURE instructors used self-efficacy talk less than four times per hour, compared to ∼60% of CURE instructors. However, this difference was not significant (*p*=0.068).

### Task Values Talk

We identified four types of task values talk, which emphasized the relevance or worth of students’ work or the course in general: *being explicit about the nature of science*, *fostering wonder*, *providing career support*, and *articulating broad relevance* (Table 5). Task values talk comprised a total of 627 quotes or 31% of the talk corpus.

**Table 5.** Task Values talk. We identified four forms of talk that emphasized the relevance or worth of students’ work or the course in general. Each form is defined in order of prevalence in the data and accompanied by example quotes. Collectively, task value related talk comprised 627 quotes, or 31% of the corpus.

| Categories | Example Quotes |
| --- | --- |
| **Being explicit about the nature of science:** Instructor comments on what science is and how science is done, including how scientists think and go about their work. (n=239; 12%) | • “Maybe a half of science is actually doing the research. Another good portion of the science is communicating your research, putting it in journal articles so that other people can read about it and presenting it in public sometimes.”<br>• “I think the results that we found were pretty exciting. There's more questions to be answered, but that's science. You start off with a question and that leads you down a different direction and it tells you what else can we do to improve the study.” |
| **Fostering wonder:** Instructor expresses interest in or enthusiasm about science or students' work. (n=206; 10%) | • “This lab is the epitome of the television description of all things in science, because it's going to be like fuchsia and blue and green and purple. All these different colors are going to come out and they're real life. They mean something and they're not just for TV. I always get excited when we have this lab.”<br>• “I'm so excited about the projects that I have lined up for this semester, so I just wanted to tease them a little bit. I've got too many projects that I want to do. A lot of great collaborations fell into place over break. Really, really cool stuff.” |
| **Providing career support:** Instructor addresses the career, academic, and/or technical needs of students. (n=99; 5%) | • “The other thing that's really interesting and important about this lab is that you are going to become much better at working with your hands. Being a nurse, you have to really become ambidextrous. You have to be able to use both hands very well to be able to manage lots of different fine motor skill tasks.”<br>• “The reason for this is if you apply for an undergraduate fellowship at a lab or a summer research opportunity, an internship, something like that, it would be nice on your resume to say things like, I am familiar with a spectrophotometer, right?” |
| **Articulating broad relevance:** Instructor comments on the larger purpose of student's work, such as how their work will be useful and used outside the classroom. (n=62; 3%) | • “So, today, we're going to look and see what your assay told us about how your enzymes work... You've contributed to a database that is going to be used by professionals from around the world, to help them improve their algorithms, to make those predictions stronger.”<br>• “This is all to say that you are all contributing to real science, that we don't have an answer to yet.” |

Instructors implicitly expressed value of the course material or activities when they were *explicit about the nature of science.* This type of talk made transparent the connections between the work students were conducting and the practices of professional scientists. For instance, one instructor explained that the class was “going to practice making observations” because “scientific pursuits begin with some observations leading to some questions.” These connections may help students see the utility value of what they were doing in class for their further education or future careers.

Instructors implied students’ work had value when they expressed their own excitement about what students were doing, the results they found, or science in general, potentially *fostering wonder* among their students as well. For example, one instructor referred to cell division as their “favorite topic” and explained, “I love it so much that you’ll get tired of me being really enthusiastic about it, but you’ll probably also be nerding out by that point, too.” By expressing their own enthusiasm or interest, instructors may spark their students’ enthusiasm about or interest in the work (i.e., intrinsic value).

Instructors made the utility value of courses or activities transparent when they *provided career support*. Career support talk involved explaining how the skills students were developing could be useful in their academic or career pursuits. For instance, one instructor expressed how the communication skills students were developing in class would be useful in a healthcare career:

Especially you guys that are pre-med, being able to actually explain to somebody who is not a scientist what something means is a skill that will serve you well in the future. You may be down the road working at a doctor’s office, and you want to explain a certain condition to your patients.

Finally, instructors made the scientific or communal value of students’ work transparent when they *articulated its broad relevance.* This type of talk emphasized the significance of students’ data or how it would be useful to the scientific community or impactful for other communities outside of the classroom. For example, one instructor contextualized students’ work by explaining how it was relevant to a community issue when they explained that:

The Forest Service put out this statement in 2014 to say that they suspect that the ground layer or the understory is in very poor condition. That’s why we need to focus on it. The restoration of that understory is underdeveloped and we don’t know how to restore it successfully, so that’s where all of you come in.

Although this quote includes references to biological content (e.g., ground layer, understory, restoration), the instructor is not defining or explaining these terms but emphasizing students’ contributions have scientific value to external stakeholders (i.e., the Forest Service).

Similar to the pattern observed for self-efficacy talk, CURE instructors used more task values talk (Median = 2.2 occurrences per hour) than non-CURE instructors (Median = 1.3 occurrences per hour) (Figure 3A). A Mann-Whitney U test comparing CUREs (n=24) and non-CUREs (n=24) showed a medium effect (Gignac & Szodorai, 2016; Hemphill, 2003) of course type on task values talk (U=505.5, r = 0.256, CI [0.02, 0.50], *p* = 0.091). Also similar to self-efficacy talk, visual inspection of the results of a KS test revealed the greatest difference between CURE and non-CURE instructors’ use of task values talk at the high end of the distribution (Figure 3B). Almost all of the instructors in our study used at least one instance of task values talk per hour regardless of their course type. However, a greater proportion of CURE instructors used more task values talk per hour, with at least half the CURE instructors in our sample using >2 instances per hour and 25% of CURE instructors using 4+ instances per hour. In contrast, ∼25% of non-CURE instructors in our sample were observed using >2 instances per hour and only one non-CURE instructor was observed using 4 instances per hour. The KS statistic (D(48)= 0.417) at ∼2.5 instances per hour supports this observation, and this difference was statistically significant (*p*=0.031).

**Figure 3.**
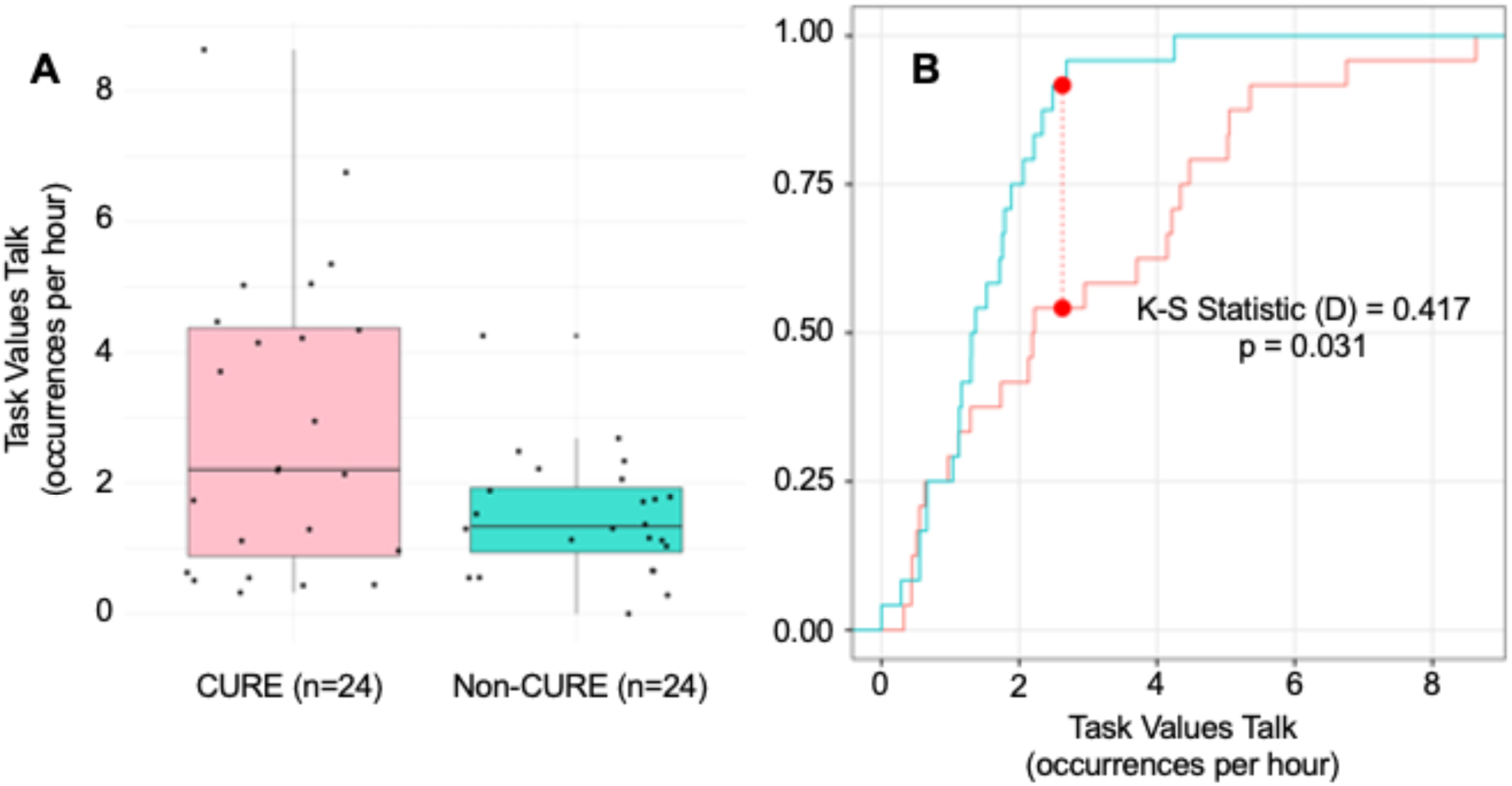
Task values talk in CUREs vs. non-CUREs. We compared occurrences of task values talk per hour for CURE instructors (coral) and non-CURE instructors (turquoise), finding that CURE instructors used more task values talk per hour than non-CURE instructors (U=505.5, r = 0.256, CI [0.02, 0.50], *p* = 0.091) (A). We also compared the distribution of task values talk between CURE and non-CURE instructors via a Kolmogorov-Smirnov test (B). We found that the point of greatest difference between the cumulative distribution functions (D(48)= 0.417) is located between two and three occurrences per hour. ∼90% of non-CURE instructors used immediacy talk less than three times per hour, compared to ∼50% of CURE instructors. This difference was significant (*p*=0.031).

We fit linear mixed effects models to independently assess whether the courses taught by self-identified CURE instructors had features typically associated with CURE instruction. Course type significantly and positively predicted students’ ratings for discovery, iteration, and cognitive ownership, although the effects were modest (standardized β=0.12, *p*= 0.0003 for discovery, β=0.09, *p*=0.0011 for iteration, and β=0.08, *p*=0.0063 for cognitive ownership; Table 6). These results indicate that, although students in CUREs reported experiencing more of these course design features than students in non-CUREs, these features vary widely within both CUREs and non-CUREs (Figure 4).

**Figure 4.**
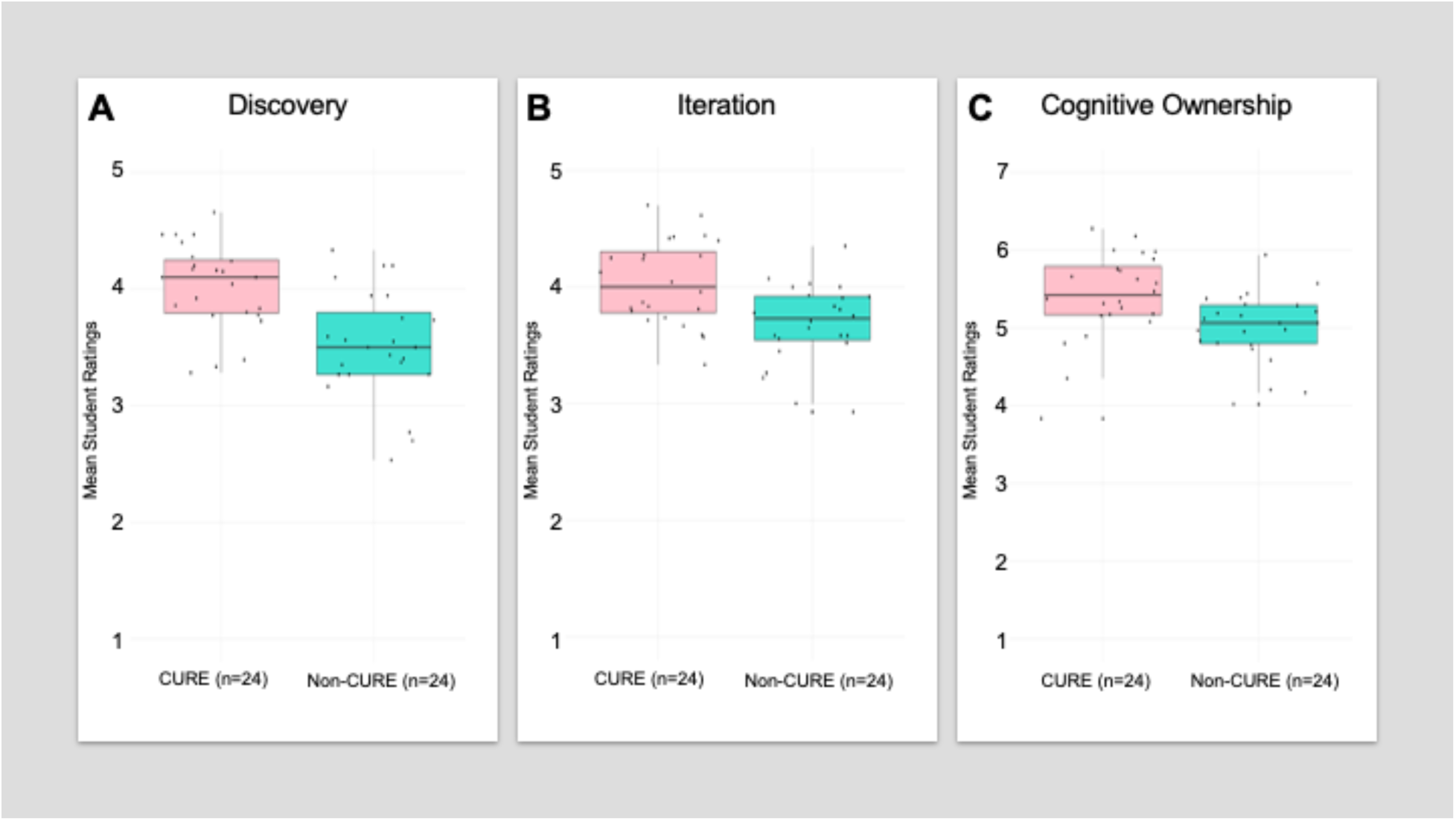
Student ratings of discovery, iteration, and cognitive ownership in CUREs vs non-CUREs. We compared student ratings of course design features typically associated with CURE instruction. Students rated discovery (A) and iteration (B) on a 5-point scale (range of 1-5) and cognitive ownership (C) on a 7-point scale (range of 1-7). Although results of our regression analysis showed that course type significantly and positively predicted students’ ratings for discovery, iteration, and cognitive ownership (see Table 6), these features vary widely within CUREs (coral) and can also be experienced by students in non-CURE courses (turquoise).

**Table 6.** Lab course design features. CURE students rated their courses more positively for discovery, iteration, and cognitive ownership than non-CURE students. Raw score means and standard deviations and standardized β are presented. Results of linear mixed models for each course design feature indicated the effect of course type on the corresponding design feature was significant, but modest.

|  | **CURE** |  | **Non-CURE** |  | **Effect of course type** |  |  |  |
| --- | --- | --- | --- | --- | --- | --- | --- | --- |
| | Mean | SD | Mean | SD | $\beta$ [CI] | SE | t | p |
| Discovery | 3.98 | 0.79 | 3.52 | 0.92 | 0.12 [0.06, 0.18] | 0.03 | t(47.0)= 3.96 | 0.0003 |
| Iteration | 4.08 | 0.68 | 3.69 | 0.82 | 0.09 [0.04, 0.14] | 0.03 | t(46.7)= 3.48 | 0.0011 |
| Cognitive Ownership | 5.50 | 0.98 | 5.05 | 1.07 | 0.08 [0.02, 0.13] | 0.03 | t(45.2)= 2.87 | 0.0063 |

## LIMITATIONS

Our study has multiple limitations that are important to consider in evaluating our results and their implications. The instructors who volunteered to participate in the study may be more student centered and thus more likely to engage in non-content instructor talk than typical lab course instructors. The instructors in our study also chose which class sessions to record, which may lead to over-representation of the forms of talk we observed. Instructors recorded only a subset of their class sessions, which may have missed important or influential forms of talk. This limitation is mitigated somewhat by the fact that many of the lab courses in our sample met once per week. Thus, recordings captured a larger proportion of the course than has been accomplished in other studies (3-4 class sessions out of a total of 12-13 weeks of class, reflecting 25-33% of the course). Some instructor and student participants did not complete the study, which could lead to bias in the results. Our sample size was small, and we conducted multiple tests, which increases risk of Type I errors. Furthermore, the small sample size limits the power of our statistical tests, which increases risk of Type II errors. Given our small sample, we opted to interpret our results with respect to effect sizes and report *p* values to allow readers to make independent judgments of any significant effects or their absence. Finally, to increase our statistical power, we treated course type as a binary rather than examining talk frequency in terms of the number of weeks students engaged in research. Below we discuss how this limitation could be addressed in future research.

## DISCUSSION

In this study, we characterized and compared the non-content talk used by CURE vs. non-CURE lab course instructors using motivation-related theoretical perspectives. We sought to explore whether differences in instructor talk had the potential to explain the motivational influence that CUREs have on students. We observed that CURE instructors used more immediacy talk than non-CURE instructors, with course type having a medium effect on immediacy talk (*p*=0.051) (Gignac & Szodorai, 2016; Hemphill, 2003). This result suggests that CUREs may on average afford more opportunities for instructors to use immediacy talk than other types of lab instruction. CURE instructors may feel they have more firsthand experiences with research than with traditional labs and thus have more fodder for sharing personal experiences with students in a CURE. Research often requires waiting, allowing for more small talk than is typical when students are conducting well-tuned lab exercises typical of traditional lab courses. Instructors may also feel like they are asking students to do something new and different in a CURE, which may prompt them to use more trust-building talk.

We also observed that CURE instructors used more self-efficacy- and task values-related talk than non-CURE instructors, with course type having a medium effect on both types of talk (Gignac & Szodorai, 2016; Hemphill, 2003). However, neither of the differences in talk met the traditional cut-off for statistical significance (i.e., *p*<0.05). CURE instructors as a group varied more widely in their self-efficacy and task values talk than non-CURE instructors, with task values talk variation meeting the traditional cut-off for statistical significance (*p*=0.031). These results suggest that CUREs may afford more opportunities for instructors to engage in self-efficacy and task values talk than non-CUREs, but that CURE instructors don’t always do so. Variation among the CUREs in our study was also evident in student ratings of discovery, iteration, and ownership, which were elevated in CUREs in our study but not distinctive to them. This result echoes a prior study showing that different types of lab instruction vary widely and often overlap in these course features (Beck et al., 2023). Considered together, these results indicate that it may be useful to study CUREs as offering varied levels of research engagement rather than as a binary of a course being a CURE or not. These results also illustrate the importance of ***measuring*** aspects of CURE design and implementation (Beck et al., 2023; Corwin, Dolan, et al., 2018; Corwin, Runyon, et al., 2015; Offerdahl et al., 2018).

We took several steps to maximize the generalizability of our results and robustness of our findings and minimize limitations. We collected data from a geographically broad sample of lab courses reflecting multiple methodological approaches (e.g., wet lab, field work, computational) taught at a diversity of institution types both on zoom and in person. We collected multiple recordings from each instructor to capture the range of talk types observable in their course. We involved multiple researchers in qualitative coding of all audio recordings to maximize consistency and minimize bias across the very large corpus. We triangulated instructor reports of course type with survey results from students. These results showed that CURE students experienced more course features associated with CURE instruction (discovery, iteration, and ownership) than non-CURE students, lending credence to our course groupings. That said, instructors in our study reported they taught CUREs when they had a research component that spanned at least one and up to 13 weeks of the course. This wide variation likely limited our ability to robustly test whether instructor talk differs between CUREs and non-CUREs.

Courses that include such disparate amounts of research are likely to be experienced quite differently by students. Prior research on CUREs demonstrates that time spent on research is an important consideration for achieving student outcomes. For example, a quasi-experimental study of one CURE program, the Freshman Research Initiative at the University of Texas Austin, showed that the full three-semester experience was necessary to observe effects on students’ graduation rates and retention in STEM majors (Rodenbusch et al., 2016). Research on another CURE program, the Genomics Education Partnership, showed that more time was necessary for students to maximize their learning (Shaffer et al., 2014). In both studies, time on research was not disaggregated from instructor talk about the lab learning experience. Time spent doing research may itself promote student learning and persistence. In addition, in more enduring research experiences, instructors may engage in more immediacy, self-efficacy, and task values talk, which then motivates student learning and persistence. Future research should aim to disentangle the effects of time spent on research and motivation-related instructor talk on the motivational effects of CUREs, either by including a consistent “dose” of research in the CURE or by collecting data from a sufficiently large and varied sample to treat research dose as a continuous variable.

Future research should examine talk longitudinally over CUREs and other lab courses to determine if talk types vary over time. Immediacy talk may be more prevalent (and potent) early in the term as instructors and students get to know one another. Self-efficacy talk may be more evident (and effective) when students encounter difficulties with coursework or failures in their investigations. Task values talk may be influential early in a course to build student buy in to research as well as during periods when research tasks become mundane because repetition is needed to address methodological issues or ensure sufficient sample sizes. Future research should also examine potential interactions between talk types. For example, immediacy talk may prompt students to pay attention during class, which promotes their learning and success and ultimately motivates them to continue in college, in science, or in research. Alternatively, immediacy talk may foster closeness such that motivational messages in the form of self-efficacy and task values talk have a stronger effect on students (i.e., immediacy talk moderates effects of self-efficacy and task values talk on student outcomes). Such research is necessary to understand whether and how instructor talk is influential for students in CUREs and other instructional contexts.

Prior research on non-content instructor talk in undergraduate courses has used a grounded approach informed by ideas of instructor immediacy, student resistance, and stereotype threat (Gelinas et al., 2022; Harrison et al., 2019; Ovid et al., 2021; Seidel et al., 2015). Our theoretical goal (i.e., talk related to motivation versus general non-content talk) and instructional context (i.e., talk in lab versus lecture courses) necessitated the development of a novel coding scheme rather than applying the existing instructor talk framework. Yet, we observed several types of talk that aligned well with the existing framework. For example, instructors in our study shared personal experiences and were explicit about the nature of science, both of which were described in the existing framework. We also identified novel types of talk that may be distinctive to lab courses or CURE instruction. For example, we observed instructors engaging in small talk, which can further build relationships. We observed instructors providing career support, encouraging students to rely on each other (i.e., interdependence), and encouraging students to take intellectual responsibility for their lab work (i.e., independence and ownership). These types of talk are consistent with elements of classroom research mentoring (Cooper & Bolger, 2024), which involves positioning students as epistemic agents to engage productively in research. Using theory, we were also able to categorize types of talk in novel ways. For example, we included sharing personal experiences in immediacy talk (versus as a stand-alone category as in the existing framework) because theory posits that disclosure helps to strengthen relationships. Informed by theory of self-efficacy and its sources, we framed self-efficacy talk as a stand-alone category rather than in terms of instructor/student relationship building. This recategorization facilitated more elaborate description of self-efficacy talk, including normalizing struggles, normalizing iteration, and providing encouragement. All of these forms of talk likely support students in navigating the failure and frustration inherent to research, which is another element of classroom research mentoring (Cooper & Bolger, 2024).

Scholars of instructor talk may wonder whether to use the existing instructor talk framework (Harrison et al., 2019; Seidel et al., 2015) or our coding scheme. Our additional characterization and refined categorization lay the groundwork to examine causal links between instructor talk and specific, motivation-related student experiences and outcomes. For example, immediacy talk should drive growth in student perceptions of closeness with their instructor, self-efficacy talk should foster students’ scientific self-efficacy, and task values talk should enhance students’ beliefs about the value of research tasks.

Which framework to use depends on the purpose and context of the research. As such, our framework is likely to be most useful for researchers interested in talk that influences motivation-related outcomes especially in the context of lab instruction.

## ACKNOWLEDGMENTS

We are grateful to our participants for allowing us to record their instruction and to their students for their survey responses. We are also grateful to Benjamin Listyg and Christina Leckfor for their guidance on data cleaning and statistical analyses. We appreciate Alexandra Cooper, Trevor Tuma, and other members of the Social Psychology of Research Experiences and Education (SPREE) research group and the Biology Education Research Group for feedback on work in progress and drafts of this manuscript. We would like to acknowledge David Esparza, Jeffrey Olimpo, and Karen Santillan for early discussions about this work. This work was supported in part by National Science Foundation Division of Undergraduate Education Improving Undergraduate STEM Education Collaborative Grant (Awards 2021138 and 2021112), an NSF Graduate Research Fellowship (Award 2236869), and the Georgia Athletic Association Professorship for Innovative Science Education. Any opinions, findings, conclusions, or recommendations expressed in this material are those of the authors and do not necessarily reflect the views of any of the funding organizations.

