## Supplemental materials for "Could instructor talk drive CURE effectiveness? A comparative study of instructor talk in introductory lab courses"

This supplement contains the following:


| Item | Page |
|---|---|
| Scales used to measure course features | S2 |
| Table S1. Standardized item loadings and standard errors | S3 |
| Table S2. Measurement model fit and reliability | S3 |
| Table S3. Intraclass correlations for course features | S3 |

**Scales used to measure course features**

We collected data using the discovery and iteration scales from the Laboratory Course Assessment Survey (LCAS) (Corwin, Runyon, et al., 2015) and the cognitive ownership scale from the Project Ownership Survey (POS) (Hanauer & Dolan, 2014).

**Discovery scale.** Students were asked to think about their course and respond as honestly and carefully as they could. Response options were: 1 – Strongly disagree; 2 – Disagree; 3 – Neither agree nor disagree; 4 – Agree, 5 – Strongly agree. Students were also given the option to select “I prefer not to respond”.

In this course, I was expected to…

1. generate novel results that are unknown to the instructor and that could be of interest to the broader scientific community or others outside the class.
2. conduct an investigation to find something previously unknown to myself, other students, and the instructor.
3. formulate my own research question or hypothesis to guide an investigation.
4. develop new arguments based on data.
5. explain how my work has resulted in new scientific knowledge.

**Iteration scale.** Students were asked to think about their course and respond as honestly and carefully as they could. Response options were: 1 – Strongly disagree; 2 – Disagree; 3 – Neither agree nor disagree; 4 – Agree, 5 – Strongly agree. Students were also given the option to select “I prefer not to respond”.

In this course, I had time to…

1. change the methods of the investigation if it was not unfolding as predicted.
2. share and compare data with other students.
3. collect and analyze additional data to address new questions or further test hypotheses.
4. revise or repeat analyses based on feedback.
5. revise drafts of papers or presentations about my investigation based on feedback.
6. revise or repeat work to account for errors or fix problems.

**Cognitive ownership scale.** Students were asked to think about their course and respond as honestly and carefully as they could. Response options were: 1 – Strongly disagree; 2 – Disagree; 3 – Somewhat disagree; 4 – Neither agree nor disagree; 5 – Somewhat agree; 6 – Agree; 7 – Strongly agree. Students were also given the option to select “I prefer not to respond”.

Please indicate the degree to which you agree with the following statements…

1. My work in this course will help solve a problem in the world.
2. My work in this course is important to the scientific community.
3. I faced challenges that I managed to overcome in this course.
4. I was responsible for the scientific outcomes in this course.
5. This course gave me a sense of personal achievement.
6. I had a personal reason for choosing what I worked on in this course.
7. The question(s) I worked on in this course I worked on was/were important to me.
8. In conducting my work in this course, I actively sought advice and assistance.

**Supplemental Table 1. Standardized item loadings and standard errors**

Confirmatory factor analyses (CFAs) were conducted and modification indices were calculated using the "lavaan" package (Rosseel, 2012) in R statistical software (R Core Team, 2022). All CFAs used maximum-likelihood (ML) estimation. Aligned with previous work assessing the measurement model of the LCAS and POS scales (see supplemental materials from Corwin et al., 2018) our modification indices suggested correlating the errors for items 3 and 4 in our Discovery scale, as well as items 1 and 2 and items 6 and 7 in our Cognitive Ownership scale. Factor loadings reported here reflect the allowance of these errors to correlate. Item numbers correspond with the order they are presented in the descriptions of the measures on the previous page.

| | **Standardized Item Loadings (Standard Errors)** | | |
|---|---|---|---|
| **Subscale** | **Discovery** | **Iteration** | **Cognitive Ownership** |
| Item 1 | 0.80 (0.05) | 0.77 (0.05) | 0.68 (0.08) |
| Item 2 | 0.78 (0.05) | 0.73 (0.05) | 0.69 (0.07) |
| Item 3 | 0.54 (0.05) | 0.40 (0.04) | 0.68 (0.05) |
| Item 4 | 0.63 (0.05) | 0.68 (0.05) | 0.57 (0.06) |
| Item 5 | 0.70 (0.05) | 0.86 (0.05) | 0.78 (0.06) |
| Item 6 | - | 0.71 (0.05) | 0.60 (0.08) |
| Item 7 | - | - | 0.72 (0.07) |
| Item 8 | - | - | 0.48 (0.06) |

**Supplemental Table 2. Measurement model fit and subscale reliability**

Measurement models denoted with an asterisk (*) reflect the modifications described in the caption for Supplemental Table 1. Coefficient omega was calculated using the "psych" package (Revelle, 2025) in R statistical software (R Core Team, 2022). Omega is not reported a second time for the models, as the reliability of the items is independent of model structural fit and thus would be equivalent.

| **Subscale** | ***df*** | **Chi-square** | **CFI** | **TLI** | **RMSEA** | **SRMR** | **ω** |
|---|---|---|---|---|---|---|---|
| Discovery | 5 | 130.73 | 0.86 | 0.73 | 0.235 | 0.06 | 0.89 |
| Discovery* | 4 | 35.98 | 0.97 | 0.91 | 0.132 | 0.03 | - |
| Iteration | 9 | 31.17 | 0.97 | 0.95 | 0.074 | 0.03 | 0.85 |
| Cognitive Ownership | 20 | 231.59 | 0.86 | 0.81 | 0.152 | 0.06 | 0.90 |
| Cognitive Ownership* | 18 | 66.97 | 0.97 | 0.95 | 0.077 | 0.03 | - |

**Supplemental Table 3. Intraclass correlations for course features**

Intraclass correlation coefficients (ICCs) and their 95% confidence intervals were calculated using the "performance" package (Lüdecke et al., 2021) in R statistical software (R Core Team, 2022).

| **Course feature** | **ICC [95% CI]** |
|---|---|
| Discovery | 0.195 [0.100, 0.356] |
| Iteration | 0.160 [0.084, 0.305] |
| Cognitive Ownership | 0.163 [0.028, 0.294] |
